# Viral and bacterial ecogenomics in globally expanding red snow blooms

**DOI:** 10.1101/2023.01.07.523113

**Authors:** Adam R. Barno, Kevin Green, Forest Rohwer, Cynthia B. Silveira

**Author notes:** Correspondence: Adam R. Barno; Cynthia B. Silveira.

## Abstract

*Chlamydomonas* algal blooms give snowmelt a red color, reducing snow albedo and creating a runaway effect that accelerates snow melting. The occurrence of red snow is predicted to grow in polar and subpolar regions with increasing global temperatures. Here, a global metaanalysis of microbial abundances showed that red snow had higher microbial abundances than white snow. We hypothesize that the increased microbial densities affect virus-bacteria interactions in snow, with potential effects on snowmelt dynamics. A genomic analysis of viral and bacterial communities in red and white snow from the Whistler region of British Columbia, Canada, identified 792 putative viruses infecting bacteria. The most abundant putative snow viruses displayed low genomic similarity with known viruses. We recovered the complete circular genomes of nine putative viruses, two of which were classified as temperate. Genomes of putative snow viruses encoded genes involved in energy metabolisms, such as NAD^+^ synthesis and salvage pathways. In model phages, these genes facilitate increased viral particle production and lysis rates. Yet, the frequency of temperate phages was positively correlated with microbial abundance in the snow samples. These results suggest the increased significance of temperate virus-bacteria interactions as microbial densities increase during snowmelt. We propose that this virus-bacteria dynamic may facilitate the red snow algae growth stimulated by bacteria, representing a potential biological feedback on snowmelt rates.

## Introduction

As the snow melts in glaciers and mountain regions, a green alga from the genus *Chlamydomonas,* which produces the red pigment astaxanthin, blooms (Duval et al., 1999; Fujii et al., 2010; Lutz et al., 2016). Astaxanthin protects the algae from UV radiation, lowers the surface albedo, and helps to warm the surrounding snow, creating a positive feedback loop of algal growth and melting snow (Fujii et al., 2010). Melting snow leads to more algae photosynthesis and labile organic carbon release, which creates a niche for nearby snow-residing heterotrophic microbes that consume the algae photosynthate (Fujii et al., 2010; Weiss, 1983) and attach to its mucilaginous coat (Weiss, 1983). Airborne microbes that accrue on the snow surface are the primary source of snow microbial communities (Harding et al., 2011; Hauptmann et al., 2014). The bacteria in snow and glacial systems display unique cryophilic features, such as an increased amount of unsaturated fatty acids and carotenoids, which increase cell membrane fluidity and stability at low temperatures (Simon et al., 2009). Snow bacteria have the highest growth rates during snow melting and have a positive influence on the growth of snow algae (Fujii et al., 2010; Harding et al., 2011; Lopatina et al., 2013; Maccario et al., 2014; Michaud et al., 2014; Segawa et al., 2005). Snow algal growth is higher in the presence of snow bacteria when compared to antibiotic-treated snow (Terashima et al., 2017; Krug et al., 2020). These observations indicate a mutualistic relationship between snow algae and associated bacteria (Yakimovich et al., 2020). The red snow algal bloom accelerates bacterial growth and changes the taxonomic and functional structure of the bacterial community (Segawa et al., 2005; Krug et al., 2020; Yakimovich et al., 2020; Hisakawa et al., 2015).

Bacteriophages (phages), viruses that infect bacteria, significantly affect bacterial population dynamics, ecology, and evolution in all aquatic, terrestrial, and animal-associated microbiomes (Breitbart et al., 2018). Yet, no study has investigated the potential role of viruses on the ecology and genomics of snow microbial communities. Phage top-down control of bacteria through virulent infection increases rates of organic matter turnover and nutrient remineralization (Breitbart et al., 2018). Therefore, virulent infections are the best-understood type of virus-bacteria ecological interactions. Yet, half of all known viruses are temperate, capable of lysogenic infections when they integrate into their host’s genome as prophages (McNair et al., 2012). During lysogenic infections, phages and their hosts establish a commensal or mutualistic relationship, with distinct outcomes for host ecology. Lysogeny may increase competitive fitness through protection against further virulent infections and frequent gene transfers through transduction (Anthenelli et al., 2020; Canchaya et al., 2003; Bondy-Denomy & Davidson, 2014). These observations from other ecosystems lead to the proposal that snow viruses also affect microbial dynamics and potentially, red snow blooms (Hisakawa et al., 2015; Rohwer et al., 2000).

Phage-encoded auxiliary metabolic genes (AMGs) modify the metabolic pathways of their microbial hosts during infection (Breitbart et al., 2007; Thompson et al., 2011). Viruses encode AMGs that complement bacterial metabolisms, such as the acquisition of alternative energy sources under starvation and UV stress protection (Breitbart et al., 2007; Puxty et al., 2018; Zeng & Chisholm, 2012). Both scenarios are relevant in the snow environment (Thomas & Duval, 1995). Hisakawa et al. (2015) found that microbial functional pathways enriched in white snow were related to glycan biosynthesis, with relatively fewer signatures of primary productivity than in red snow. Microbial functional pathways enriched in red snow were related to photosynthesis, carbon fixation, and methane metabolism (Hisakawa et al., 2015).

Here, we investigated viral and bacterial ecology and genomics in red and white snow through a meta-analysis of global samples and genomics from three locations in British Columbia, Canada. Global abundance data showed increased microbial densities in red snow, while the samples analyzed here for viral abundance displayed low virus-to-microbe ratios. The dominant putative snow viruses were virulent phages genomically distinct from known viruses. Snow virus genomes encode auxiliary metabolic genes (AMGs) involved in energy production and protection against host defense, which are typically involved in virulent infection. These results suggest that lysis-lysogenic switches take place during red snow bloom, potentially affecting bloom progression.

## Methods

### Meta-analysis of viral and bacterial abundances

Snow bacterial cell abundances were collected from nine studies listed in Table 1 (Amato et al., 2007; Carpenter et al., 2000; Harding et al., 2011; Liu et al., 2009; Lopatina et al., 2013; Lutz et al., 2016; Michaud et al., 2014; Segawa et al., 2005; Thomas & Duval, 1995). These abundances were combined with new abundance data obtained from the Whistler region of British Columbia, Canada (described below). Log-transformed cell abundances in red snow and white snow were normally distributed (p > 0.05, Shapiro – Wilk test) and were compared between snow types using a t-test.

**Table 1.** Snow metadata.

| Sample Site | Snow type | Elevation (m) | Location | Cells ($10^5$ / mL) | Shannon ($H'$) | Reference |
| --- | --- | --- | --- | --- | --- | --- |
| BCWS | white | 1831 | Canada | 16 | 3.14 | Present Study |
| BCRS | red | 1826 | Canada | 21 | 2.79 | Present Study |
| CAWS | white | 847 | Canada | 50 | 2.96 | Present Study |
| CARS | red | 829 | Canada | 49 | 2.83 | Present Study |
| CGRS | red | 989 | Canada | 67 | 2.85 | Present Study |
| Amundsen-Scott South Pole<br>Station | white | 2835 | Antarctica | 0.031 | N/A | Carpenter et al., 2000 |
| Druzhnaja, 54th | white | 3 | Antarctica | 0.46 | 2.34 | Lopatina et al., 2013 |
| Leningradskaja, 54th | white | 300 | Antarctica | 0.37 | 2.49 | Lopatina et al., 2013 |
| Druzhnaja, 55th | white | 3 | Antarctica | 0.056 | 1.75 | Lopatina et al., 2013 |
| Leningradskaja, 55th | white | 300 | Antarctica | 0.010 | 2.26 | Lopatina et al., 2013 |
| Concordia | white | 3233 | Antarctica | 0.0043 | N/A | Michaud et al., 2014 |
| Svalbard | red | 274 | Arctic | 0.44 | 5.67 | Lutz et al., 2016 |
| Tarfala,<br>Northern<br>Sweden | red | 1264 | Arctic | 0.12 | 4.95 | Lutz et al., 2016 |
| Site E | red | 3050 | California | 6.0 | N/A | Thomas and Duval, 1995 |
| Site F | red | 3050 | California | 3.6 | N/A | Thomas and Duval, 1995 |
| Site G | red | 3050 | California | 3.4 | N/A | Thomas and Duval, 1995 |
| Site E | white | 3050 | California | 1.6 | N/A | Thomas and Duval, 1995 |
| Site F | white | 3050 | California | 1.3 | N/A | Thomas and Duval, 1995 |
| Site G | white | 3050 | California | 0.40 | N/A | Thomas and Duval, 1995 |
| Ellesmere Island | white | 0 | Canadian Arctic | 0.27 | 1.65 | Harding et al., 2011 |
| Char Lake | white | N/A | Canadian Arctic | 0.011 | 0.36 | Harding et al., 2011 |
| Murodo-daira | white | 2450 | Japan | 0.84 | N/A | Segawa et al., 2005 |
| Kuranosuke snow patch | white | 2600 | Japan | 0.84 | N/A | Segawa et al., 2005 |
| Hamaguri snow patch | white | 2700 | Japan | 0.84 | N/A | Segawa et al., 2005 |
| Ny-Ålesund | white | 10 | Svalbard | 0.60 | N/A | Amato et al., 2007 |
| Kongsvegen glacier | white | 670 | Svalbard | 0.25 | N/A | Amato et al., 2007 |
| Guoqu Glacier | white | 6621 | Tibetan Plateau | 0.35 | 2.5 | Liu et al., 2009 |
| Zadang Glacier | white | 5799 | Tibetan Plateau | 5.4 | 4.0 | Liu et al., 2009 |
| East Rongbuk Glacier | white | 6520 | Tibetan Plateau | 0.12 | 2.2 | Liu et al., 2009 |
| Palong No. 4 Glacier | white | 5507 | Tibetan Plateau | 0.11 | 3.1 | Liu et al., 2009 |

N/A: Data not available.

### Sample Collection

Snow samples (30 L) were collected in June 2017 in the Whistler region of British Columbia, Canada. Five samples were collected from three different locations using a sterile shovel and 30L bags: Blackcomb Mountain (BC) on the Catskinner Ski Run (red snow: BCRS, 50° 5’ 44.8188’’ N, 122° 54’ 13.4388’’ W; white snow: BCWS, 50° 5’ 43.0188’’ N, 122° 54’13.2012’’ W), adjacent to Callaghan Creek (CA) near the Callaghan Ski Run (red snow: CARS, 50° 8’ 59.19’’ N, 123° 7’ 18.4188’’ W; white snow: CAWS, 50° 8’ 59.1612’’ N, 123° 7’ 13.8’’ W), and on the Sixteen Mile Creek Forest Service Road on Cougar Mountain (CG, red snow: CGRS, 50° 10’ 50.304’’ N, 122° 58’ 25.0356’’ W) (Figure 1A, inset; Table 1). Within two hours of collection, snow samples were melted at room temperature and filtered through a 25 μm Nitex mesh filter to remove large debris.

**Figure 1.**
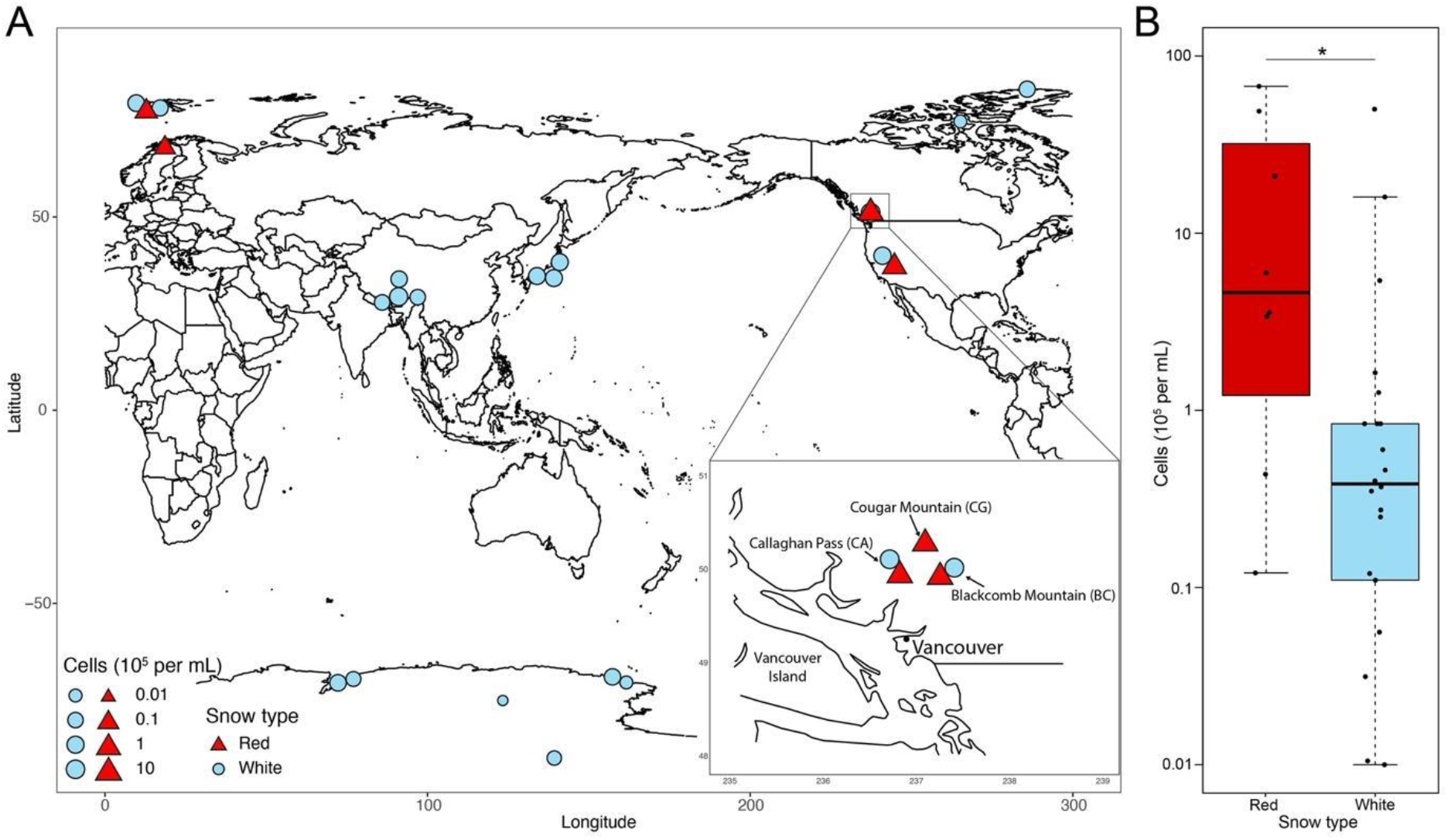
Microbial abundances in red and white snow. (A) Locations where microbial cell abundances have been quantified in snow samples including the present study (inset) and nine previous studies (Table 1). (B) Boxplot displaying the cell abundances in red snow (n = 8) and white snow (n = 22), (*t* (12.9) = 2.9, *p* = 0.009). Map lines do not necessarily depict accepted national boundaries.

### Bacterial and viral abundances from the Whistler region

For bacterial abundance determination in the Whistler region, 1 mL from the 25 μm-filtered snow samples was fixed with 2 % paraformaldehyde. These subsamples were passed through a 0.02 μm Anodisc filter (Whatman, UK), and the particles remaining on the filter were stained with SYBR Gold (Life Technologies, USA), mounted on slides, and analyzed by epifluorescence microscopy (Thurber et al., 2009). Ten random images were obtained using 600X magnification from each snow sample, and bacteria and viruses were counted using Image-Pro Plus (Media Cybernetics, MD). The total number of viruses and bacteria per mL of snow was then calculated using the numbers obtained from the microscopy images corrected for the area of each field of view. Virus-to-microbe ratios obtained from the snow samples were compared with published VMRs from 23 studies covering eleven ecosystems (Anthenelli et al., 2020).

### DNA extraction and sequencing

The remaining 25 μm-filtered snowmelt (~30L) was concentrated using a 100 kDa tangential flow filtration system to a final volume of 500 mL. The concentrate was filtered through a Sterivex 0.45 μm PVDF membrane filter. Chloroform was added to the filtrate at a final concentration of 0.5 % and stored at 4 °C until further purification of the viral fraction in the laboratory. The membrane filter was stored at −20 °C until processing for DNA extraction.

Viral concentrates were further concentrated in the laboratory using 10 % (w/v) polyethylene glycol (PEG) 8000. After overnight incubation, the concentrates were centrifuged at 11,000 × g for 40 minutes at 4 °C. Supernatants were removed, and the concentrated pellets were resuspended in 8 mL of Saline Magnesium buffer (Lim et al., 2014), which was then overlayed onto a Cesium Chloride gradient containing 1 mL each of 0.02 μm-filtered Cesium Chloride – SM buffer solutions at successive densities of 1.2 g/mL, 1.35 g/mL, 1.5 g/mL, and 1.7 g/mL and centrifuged at 82,844 × g (22,000 RPM) for 12 hours at 4 °C (Lim et al., 2014). After centrifugation, 1 mL of the 1.5 g/mL fraction containing the putative purified viral fraction was extracted. The purity of the viral fraction was verified by visualizing 100 μL of each sample stained with SYBRGold on an epifluorescence microscope. DNA was extracted from the concentrated viral particles following the formamide/cetyltrimethylammonium bromide (CTAB) and phenol/chloroform methods outlined in Thurber et al., 2009. Extracted viral DNA was resuspended in 50 μL of molecular-grade water. Removal of microbial DNA was verified using 16S rDNA PCR with the primers 27F (5’ - AGR GTT TGA TCM TGG CTC AG - 3’) and 1492R (5’ - GGH TAC CTT GTT ACG ACT T - 3’). Temperature cycling for the PCR was 95 °C for 5 min, followed by 30 cycles of 95 °C for 0.5 min, 51 °C for 0.5 min, and 72 °C for 2 min, with one final cycle of 72 °C for 10 min. Only samples with no 16S amplification in the PCR were kept for further steps. DNA libraries for shotgun metagenomic sequencing of viral concentrates were prepared using an Accel-NGS^®^ 1S Plus DNA Library Kit (Swift Biosciences, USA) and sequenced on an Illumina MiSeq platform (Illumina, USA) using Illumina MiSeq Reagent Kit V3 (Illumina, USA). Reads were quality-controlled using BBduk for adaptor trimming, quality trimming (>30), quality filtering (>30), K-mer filtering (which removes reads with a 31-mer match to an Illumina spike-in), and Entropy filtering (>0.90) (Bushnell, 2014). Reads were aligned to the human reference genome GRCh38 database using Smalt with an 80% identity threshold to remove potential contaminants; however, no sequences were removed in this step (Ponstingl & Ning, 2010). The number of reads produced from the red snow virome samples was three times higher than that produced from the white snow samples (91,254,046 reads to 29,902,344 reads, respectively). The number of hits to the snow virome database also differed between red snow, with 2,385,982 total hits or 97 %, and white snow, with 64,999 total hits or 2.7 %.

Microbial metagenomic DNA was extracted by inserting the 0.45 μm filters in the PowerSoil DNA Isolation Kit (Qiagen, Germany) bead tubes and following the manufacturer’s instructions. Metagenomic DNA libraries were prepared using KAPA Hyper P Kits (Roche, Switzerland), and each sample was sequenced in 3 different lanes (3 technical replicates per sample) on an Illumina HiSeq platform (Illumina, USA).

### Taxonomic and functional assignments of bacterial communities

Raw metagenomic reads from the 0.45 μm filter obtained from the five sites were uploaded to the MG-RAST server (Meyer et al., 2008) and quality-controlled by adaptor trimming, quality trimming (>30), and quality filtering (>30). Sequences were compared to the M5NR database for taxonomic assignment requiring a minimum length of 15 amino acids with at least 60 % identity, and an e-value less than 10^-5^. Relative abundances were calculated by taking the number of hits per taxon divided by the total number of hits. For the functional annotation, sequences were compared to the SEED database with a minimum length of 15 amino acids with at least 60 % identity, and an e-value less than 10^-5^ (Meyer et al., 2008).

### De novo virome analysis

Viromes from the five Canada snow samples were assembled into contigs using metaSPAdes (Nurk et al., 2017). Contigs from each individually assembled sample were combined and clustered by 98% similarity using CD-HIT (Li & Godzik, 2006). Contigs longer than 1 Kb were submitted to VIBRANT (Virus Identification by IteRative ANnoTation) phage annotation tool (Kieft et al., 2020). VIBRANT uses protein annotations from Hidden Markov Models (HMMs) and a “v-score” metric to identify viral sequences (Kieft et al., 2020). 792 unique viral contigs identified by VIBRANT composed the snow virome database. Post-QC virome and metagenome fastq reads were mapped to this database using Smalt with an 80% identity threshold (Ponstingl & Ning, 2010). A nonredundant list of viral contigs recruiting more than one read was generated from mapping results, which resulted in 792 unique contigs. The percent abundance of each viral contig per sample was calculated and sorted to select the ten most abundant contigs per sample, for a total of 36 unique contigs across all samples.

Phylogenomic analysis of the top 36 viral contigs was performed using the ViPTree server (version 1.9), which produced a viral proteomic tree with closely related reference viral genomes from NCBI RefSeq and GenBank (Nishimura et al., 2017). The tree was generated using a protein distance metric based on similarities calculated from normalized tBLASTx scores (Nishimura et al., 2017). The 36 viral contigs were compared with the 8 most closely related genomes (from genome similarity S_G_ scores) for each contig. The tree included 131 dsDNA phages and the 36 viral contigs obtained in this study.

Read recruitment by the most abundant contigs was visualized using the Anvi’o metagenomic workflow in Anvi’o-6.2 (Eren et al., 2015). Functional annotations of viral contigs were determined via Hidden Markov Model (HMM) searches against the Pfam, KEGG, and VOG databases within VIBRANT. VIBRANT also identified complete circular phages, temperate phages (due to the presence of an integrase gene), and phage-encoded Auxiliary Metabolic Genes (AMGs) (Kieft et al., 2020). The correlation between the percent abundance of temperate phages and cell abundance counts was tested using the Spearman correlation in the *psych* R package (Revelle, 2021). Metabolic pathway affiliations of the AMGs were derived from the KEGG pathway database (Kanehisa & Goto, 2000). AMG abundance was determined using read recruitment for each AMG-containing contig. The six complete circular phages containing at least one AMG were visualized using DNAplotter (Carver et al., 2009).

## Results

### Microbial abundance and community composition

A meta-analysis of abundance data from the present study and nine previous studies of globally distributed sites showed that microbial abundances in red snow were nearly fivefold higher than those in white snow (*t* (12.9) = 2.9, *p* = 0.013, average microbial abundances ± SEM: 1.9 ± 0.9 × 10^6^ cells/mL for red snow (n = 8), 3.6 ± 2.3 × 10^5^ cells/mL for white snow (n = 22); Figure 1A and 1B, Table 1). Viral abundances for the five Canada snow samples ranged from 1.1 × 10^6^ virus-like particles per mL (VLPs/mL) in BCWS to 2.3 × 10^6^ VLPs/mL in BCRS (Table 2). VMRs were higher in the BC samples (VMR = 6.8 in BCWS and 11 in BCRS) than in the CA samples (3.9 in CAWS, 3.1 in CARS) and CGRS (2.1). VMRs in the snow samples were comparably low when plotted with VMRs observed in other ecosystems (Supplementary Figure 1).

**Table 2.** Microbial cell abundances, viral abundances, and virus-to-microbe ratios in the Canada snow samples.

| Sample | Snow type | Cells ($10^6$ / mL) | Virus-like particles ($10^6$ / mL) | Virus-to-microbe ratio |
| --- | --- | --- | --- | --- |
| BCWS | white | 1.6 | 10.9 | 6.8 |
| BCRS | red | 2.1 | 23.2 | 11 |
| CAWS | white | 5.0 | 19.5 | 3.9 |
| CARS | red | 4.9 | 15.2 | 3.1 |
| CGRS | red | 6.7 | 14.1 | 2.1 |

The percent abundance of reads belonging to the algae genus *Chlamydomonas* in the snow samples ranged from 0.1 % of the total reads in CAWS to 1.8 % in BCRS (Supplementary Figure 2). The most abundant bacterial phyla in the Canada snow metagenomes were Proteobacteria (56 % of the total bacterial reads), Bacteroidetes (30 %), Actinobacteria (7.2 %), Acidobacteria (2.2 %), Firmicutes (1.9 %), and Cyanobacteria (1.1 %) (Supplementary Figure 3A). The most abundant bacterial classes were Betaproteobacteria (28 % of the total bacterial reads), Alphaproteobacteria (14 %), Sphingobacteria (14 %), Gammaproteobacteria (12 %), and Actinobacteria (7.2 %) (Supplementary Figure 3B). Metagenomic functional profiles were assigned by classifying reads according to the SEED subsystems levels 1, 2, and 3 (Supplementary Figures 4A, B, and C, respectively). Among the eight most abundant SEED level 2 subsystems were functions related to protein biosynthesis; central carbohydrate metabolism; monosaccharides; lysine, threonine, methionine, and cysteine; and folate and pterins (Supplementary Figure 4B).

### Abundant and divergent snow viruses

Thirty-six unique viral contigs comprised the ten most abundant viral contigs in each sample. Although no viral contig was within the top 10 most abundant in all viromes, nine viral contigs were within the most abundant in two or more samples. Five of the 36 contigs were classified as temperate phages, while the other 31 were classified as virulent phages based on the presence or absence of VOG-annotated integrase genes (Kieft et al., 2020). The relationship between these 36 contigs and reference viral genomes from the GenomeNet Virus-Host DB (which contains viruses with complete genomes from NCBI RefSeq and Genbank whose accession numbers are listed in EBI Genomes) was visualized using a viral proteomic tree (Nishimura et al., 2017) (Figure 2). The snow contigs formed branches distantly related to reference genomes, indicating that these contigs represent novel viral taxa from the snow biome. The hosts of the reference viruses with the closest relationship with snow viruses were Gammaproteobacteria, Cyanobacteria, Firmicutes, Alphaproteobacteria, and Actinobacteria, reflecting the bacterial community composition in snow. Three snow virus clades with deep branching in this tree were formed, containing seven viral contigs each. Clade 1 also included two Pseudomonas phages and one Ralstonia phage. The most closely related reference genomes to Clade 2 were a Cronobacter phage, Klebsiella phage, Enterobacter phage, and two Escherichia phages. Clade 3 formed a group of distantly related viruses containing both virulent and temperate phages whose closest references were Cellulophaga phages. The read recruitment by the snow virome database demonstrated that most contigs were abundant in only one or a few samples (Figure 3). Red snow recruited reads at a more similar level among samples than the white snow metagenomes and viromes. Nine of the 36 viral contigs were determined to be putative complete circular genomes (Kieft et al., 2020). Two of the nine complete viral genomes were classified as temperate by the presence of an integrase (Kieft et al., 2020).

**Figure 2.**
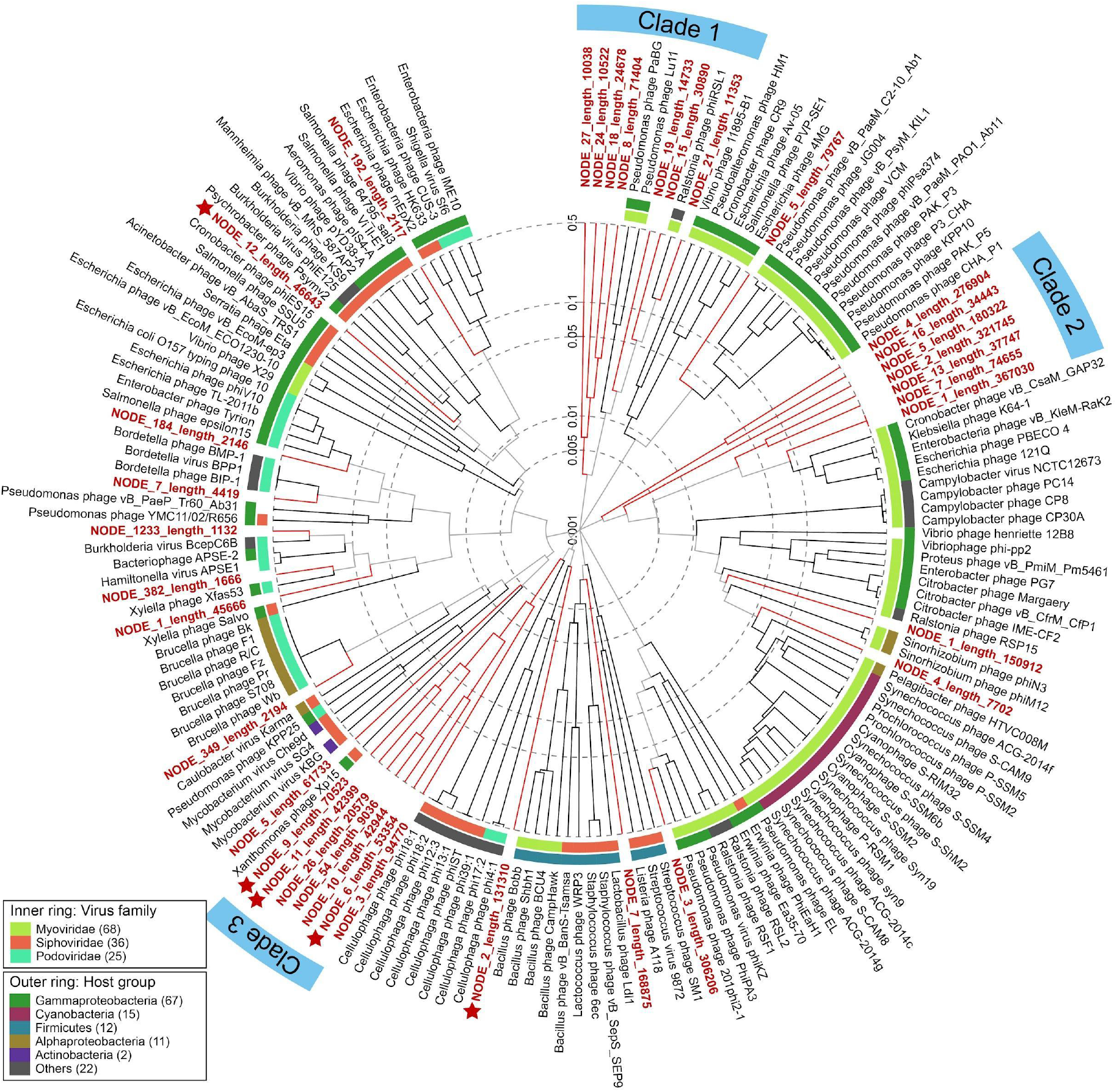
Viral proteomic tree of the ten most abundant viral contigs and their most closely related reference genomes. The host groups of the reference viruses indicate probable hosts for the newly defined snow viruses. Stars indicate temperate phages from this study. Branches are shown in Log scale.

**Figure 3.**
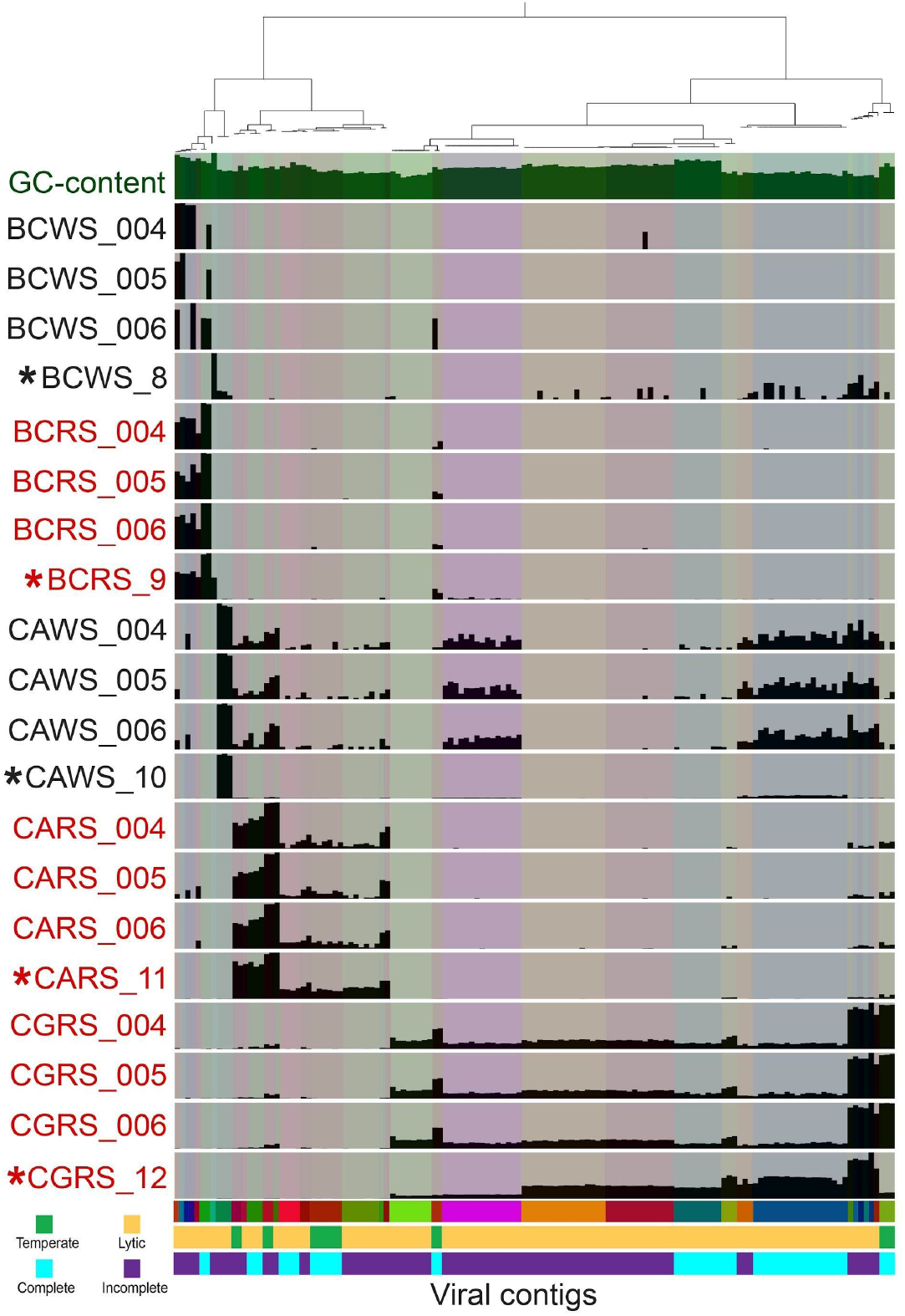
Abundances of the ten most abundant viral contigs in each of the samples (36 across all samples). Abundances were visualized using the Anvi’o metagenomic workflow. Hierarchical clustering of the contig bins is done based on Ward’s method using Euclidean distances. The heights of the bars display the mean coverage of each contig divided by mean coverage of all of the contigs. BCWS = Blackcomb Mountain White Snow, BCRS = Blackcomb Mountain Red Snow, CAWS = Callaghan Pass White Snow, CARS = Callaghan Pass Red Snow, CGRS = Cougar Mountain Red Snow. Asterisks denote virome samples.

### Prevalence of temperate phages

Of the 792 unique viral contigs, 20 were classified as being from a temperate phage (2.5 %). The relative abundance of temperate phages in the microbial fraction, determined by read recruitment, ranged from 0 % to 31 %, and the average percent abundance of temperate phages across all samples was 11 % (Table 3). The relative abundance of temperate phages in the microbial fraction was significantly correlated with cell abundance (Spearman, *r_s_*(13) = 0.66, *p* = 0.008). In the two sample sites with paired white and red snow samples (BC and CA), the percent abundances of temperate phages were higher in the red snow samples (Table 3).

**Table 3.** Abundance of temperate phages at each sample site. BCWS = Blackcomb Mountain White Snow, BCRS = Blackcomb Mountain Red Snow, CAWS = Callaghan Pass White Snow, CARS = Callaghan Pass Red Snow, CGRS = Cougar Mountain Red Snow. Asterisks denote virome samples.

| Sample | Temperate (%) | Virulent (%) |
| --- | --- | --- |
| BCWS_L004 | 0 | 100 |
| BCWS_L005 | 0 | 100 |
| BCWS_L006 | 0.391 | 99.6 |
| BCWS_S8* | 1.31 | 98.7 |
| BCRS_L004 | 0.967 | 99 |
| BCRS_L005 | 0.782 | 99.2 |
| BCRS_L006 | 1.01 | 99 |
| BCRS_S9* | 1.91 | 98.1 |
| CAWS_L004 | 11.5 | 88.5 |
| CAWS_L005 | 12 | 88 |
| CAWS_L006 | 11.4 | 88.6 |
| CAWS_S10* | 22.3 | 77.7 |
| CARS_L004 | 23.3 | 76.7 |
| CARS_L005 | 24.5 | 75.5 |
| CARS_L006 | 30.6 | 69.4 |
| CARS_S11* | 30.4 | 69.6 |
| CGRS_L004 | 22.5 | 77.5 |
| CGRS_L005 | 1.3 | 98.7 |
| CGRS_L006 | 22.3 | 77.7 |
| CGRS_S12* | 1.16 | 98.8 |
| Total | 11.0 | 89.0 |

### Auxiliary metabolic genes

The viral contigs encoded 46 unique auxiliary metabolic genes (AMGs), four in viruses classified as temperate. The most abundant AMGs across all samples were *NAMPT* (nicotinamide phosphoribosyltransferase), *nadM* (bifunctional NMN adenylyltransferase/nudix hydrolase), *folA* (dihydrofolate reductase), *nadR* (HTH-type transcriptional regulator), and *DNMT1* (DNA (cytosine-5)-methyltransferase 1). The AMGs encoded by temperate viruses were *mec* ([CysO sulfur-carrier protein]-S-L-cysteine hydrolase), *DNMT3A* (DNA (cytosine-5)-methyltransferase 3A), *moeB* (molybdopterin-synthase adenylyltransferase), and *sacA* (beta-fructofuranosidase), which are broadly involved in the cysteine biosynthesis (*mec*), DNA methylation (*DNMT3A*), carbohydrate metabolism (*sacA*), and the sulfur relay system (*moeB*).

Figure 4A and Figure 4B show the ten most abundant AMGs in the viromes and metagenomes, respectively, as determined via read recruitment by the AMG-containing contigs. Five AMGs were shared between virome and metagenome contigs: *NAMPT*, *nadM*, *folA*, *nadR*, and *queD* (6-pyruvoyltetrahydropterin/6-carboxytetrahydropterin synthase). Among the most abundant AMGs that were not shared, the viral fraction had genes related to coenzyme A biosynthesis (*coaE*; dephospho-CoA kinase, and *coaD*; pantetheine-phosphate adenylyltransferase) and NAD^+^ synthesis (*nadE*; NAD^+^ synthase), as well as *UGDH* (UDP glucose 6-dehydrogenase), *rfbA* (glucose-1-phosphate thymidylyltransferase), and *gpmB* (probable phosphoglycerate mutase). The uniquely abundant AMGs in the microbial fraction were *DNMT1* and *DNMT3A, mec, queE* (7-carboxy-7-deazaguanine synthase), and *folE* (GTP cyclohydrolase IA). Two of these AMGs (*mec* and *DNMT3A*) were found in temperate phages, as well as in virulent phages.

**Figure 4.**
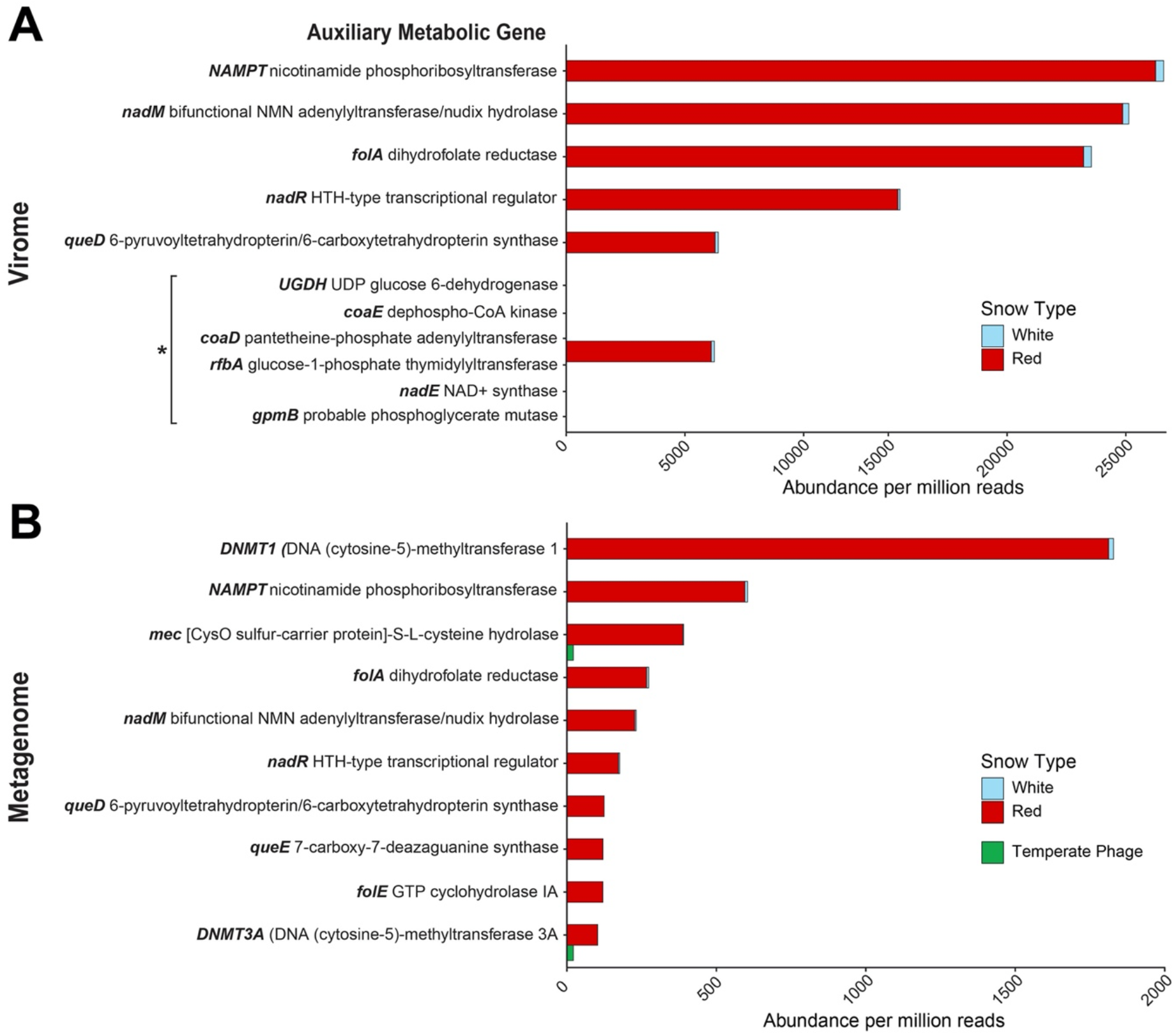
Abundance of dominant Auxiliary Metabolic Genes (AMGs) in the virome (A) and in viral sequences identified in the metagenome (B). Data are shown as the abundance of the AMG-containing contig per one million reads. Five of the most abundant AMGs were shared between the virome and the metagenome. AMGs in temperate phages were found only within the top 10 AMGs in the metagenomic fraction. *All the six AMGs indicated by the asterisk were predicted to be encoded by a single putative viral genome fragment (contig NODE_2_length_321745).

## Discussion

The effects of phage-bacteria interactions on microbial communities subject to extreme cold, desiccation, and UV exposure in snow environments are entirely unknown (Harding et al., 2011). This represents a significant knowledge gap considering the increase in global temperatures and expanding red snow events. Here, we show that snow viruses are mostly uncharacterized, forming distinct clades from all known reference viral genomes. All closely related viruses are tailed phages, which suggests that tailed phages also dominate the snow virome. The most abundant novel snow phages have predicted hosts within the class Gammaproteobacteria, the fourth most abundant class in the microbial metagenomes.

The most abundant AMGs found in the Canada snow viral genomes indicate that the viruses direct the energy of the host to produce new viral progeny and evade host defenses during virulent infection. Three genes involved in the NAD^+^ salvage pathway from nicotinamide (*NAMPT*, *nadM*, and *nadR*) were among the most abundant AMGs in both the virome and metagenome fractions. *NAMPT* is the rate-limiting step in NAD^+^ synthesis (Revollo et al., 2007), and *nadM* and *nadR* are both adenylyltransferases involved in the NAD^+^ synthesis/salvage pathways (Lau et al., 2009). Six viral contigs encoded both enzymes (*NAMPT* and *nadM)* necessary to produce NAD^+^ from nicotinamide. Each of these contigs was predicted to be from a virulent phage, including the two complete circular contigs containing both genes (NODE_1_length_367030 and NODE_5_length_180322). Another gene involved in this pathway, *nadE*, was found among the most abundant AMGs in the virome fraction. *nadE* encodes an enzyme involved with synthesizing NAD^+^ *de novo* via different intermediates, a process first discovered in *Francisella tularensis* (Sorci et al., 2009). The production of NAD^+^ salvage enzymes in Vibrio Phage KVP40 increases upon infection of its host, which suggests that NAD^+^ biosynthesis after phage infection aids in the production of viral progeny (Lee et al., 2017). In other ecosystems, NAD^+^-dependent catabolic pathways are more abundant in low-density microbial communities where virulent infection is dominant (Silveira et al., 2019; Knowles et al., 2016). This suggests that, in snow, phage-encoded AMGs for NAD^+^ metabolism contribute to virulent phage production.

Viral AMG functions are associated with the functional capacities of the microbial hosts. For example, the most abundant SEED level 3 subsystem in the metagenome, Respiratory Complex I (NADH:ubiquinone oxidoreductase), uses the energy released by the electron transfer from NADH to quinone to pump protons across the plasma membrane (Walker, 1992). The bioavailability of this complex could be modified via the expression of viral auxiliary metabolic genes involved in NAD^+^ biosynthesis. The ubiquitous presence of *nadM* in the snow viromes and 4 AMGs involved in NAD^+^ biosynthesis pathways (*NAMPT*, *nadM*, *nadR*, and *nadE*) support this hypothesis. These genes were found in virulent and temperate phages from red and white snow and indicate how host metabolisms can be modified by infecting phages to control the catabolism in their bacterial hosts.

Dihydrofolate reductase (*folA*) was another abundant gene that functions as a structural protein in phage baseplates (Kozloff et al., 1977) and as a metabolic enzyme involved in thymine synthesis (Mathews, 1967). The latter function is essential in maintaining the high rate of DNA synthesis that occurs in T4 phage-infected cells (Mathews, 1967). Similarly, two genes related to the synthesis of coenzyme A (CoA), *coaD* and *coaE*, were abundant in the virome fractions of the snow samples. CoA is a cofactor that participates in multiple energy metabolism pathways (Leonardi et al., 2005). Intermediate metabolites in the coenzyme A biosynthesis pathway have been shown to increase during virulent phage infection (De Smet et al., 2016), although the purpose of this pathway in phage lysis is not entirely known.

The AMGs *queD* (in both the virome and metagenome), as well as *queE* and *folE* (among the most abundant AMGs in the metagenome), encode enzymes involved in the synthesis of 7-cyano-7-deazaguanine from GTP in the queuosine biosynthesis pathway (Hutinet et al., 2019; Phillips et al., 2008; Reader et al., 2004). Queuosine is a modified base in the wobble position of tRNAs that increases the fidelity of protein synthesis (Vinayak & Pathak, 2010). Moreover, 7-deazaguanine derivatives produced in this pathway protect phage DNA from host restriction enzymes without affecting DNA polymerase activity (Hutinet et al., 2019; Mačková et al., 2015). Each of these genes was only found within virulent phages in the Canada snow samples. In a related mechanism, DNA methyltransferases also help phages to avoid host defenses. Two different DNA methyltransferase genes (*DNMT1* and *DNMT3A*) were abundant in the snow viral contigs. Depending on the target sequence, DNA methyltransferases control the expression of genes by adding methyl groups to CpG structures in DNA (Sánchez-Romero et al., 2015). Canonically, bacteria defend against phage infection via restriction-modification (R-M) systems consisting of restriction enzymes and methyltransferases (Tock & Dryden, 2005). By methylating their DNA at specific sites, bacteria distinguish between self DNA (methylated) and invading phage DNA (nonmethylated) and subsequently digest the invading DNA by restriction enzymes. However, certain phages avoid bacterial restriction enzymes by methylating their own DNA during virulent production or prophage induction, thereby disguising the phage DNA within the bacterial host (Hampton et al., 2020; Krüger & Bickle, 1983; Murphy et al., 2013; Samson et al., 2013). DNA methyltransferases were present in both virulent and temperate phages from the Canada snow samples. The abundance of these defense-counter-defense genes suggests that a virulent arms race occurs in snow environments.

While virulent infection appears to be the dominant viral behavior in snow environments, the combination of genomic and density data indicates changes in virus-bacteria interactions in melting snow affected by algae blooms. Microbial abundances from nine published studies with a wide geographical range showed that the abundance of microbes in red snow was higher than in white snow (Figure 1). This could be due to increased microbial growth rates in red snow, fueled by a positive feedback loop of microbial and algal growth as the algae release organic carbon that fuels heterotrophic bacteria (Hisakawa et al., 2015). High microbial density increases the frequency of phage-host encounters and, therefore, the frequency of lysogeny caused by phage coinfections (Luque & Silveira, 2020; Knowles et al., 2016). High densities fueled by the increase in labile organic carbon are also typically associated with metabolic switches toward low-efficiency metabolisms that rely more on NADPH than NADH (Silveira et al., 2021). In red snow, this switch is likely fueled by the organic carbon released by the algae. Therefore, the increase in bacterial densities in red snow leads to the hypothesis that snow communities switch from lysis to lysogeny during red snow blooms. The positive correlation between cell abundance and temperate phage relative abundance in the metagenomes of the Whistler region supports this hypothesis (Spearman, *r_s_*(13) = 0.66, *p* = 0.008). Similarly, the highest virus-to-microbe ratios (VMRs) were found in BCWS and BCRS, with the lowest relative abundances of temperate phages and cell abundances (Figures 1 and 4). These results suggest that density-dependent transitions from virulent to temperate at high microbial abundances, as predicted by the Piggyback-the-Winner dynamics, also occur in snow ecosystems. Future studies with broader geographical ranges and sample sizes are necessary to understand the biogeographical patterns associated with microbial and viral density and infection dynamics in snow.

## Conclusion

Viruses play essential roles in biogeochemical cycling and microbial ecology in all the studied planet’s biomes. Here, we showed that viruses are active members of the snow microbial community, with potential implications for snowmelt dynamics. Our data suggest that viruses control bacterial populations through lytic induction and redirect the host’s energetic metabolism, likely impacting organic matter turnover. The increase in microbial abundances in red snow blooms is associated with increasing temperate viral behavior. This trend is predicted to decrease viral top-down control on bacteria, accelerating the bacteria-algae feedback loop that facilitates snowmelt and decreases albedo.

**Figure 5.**
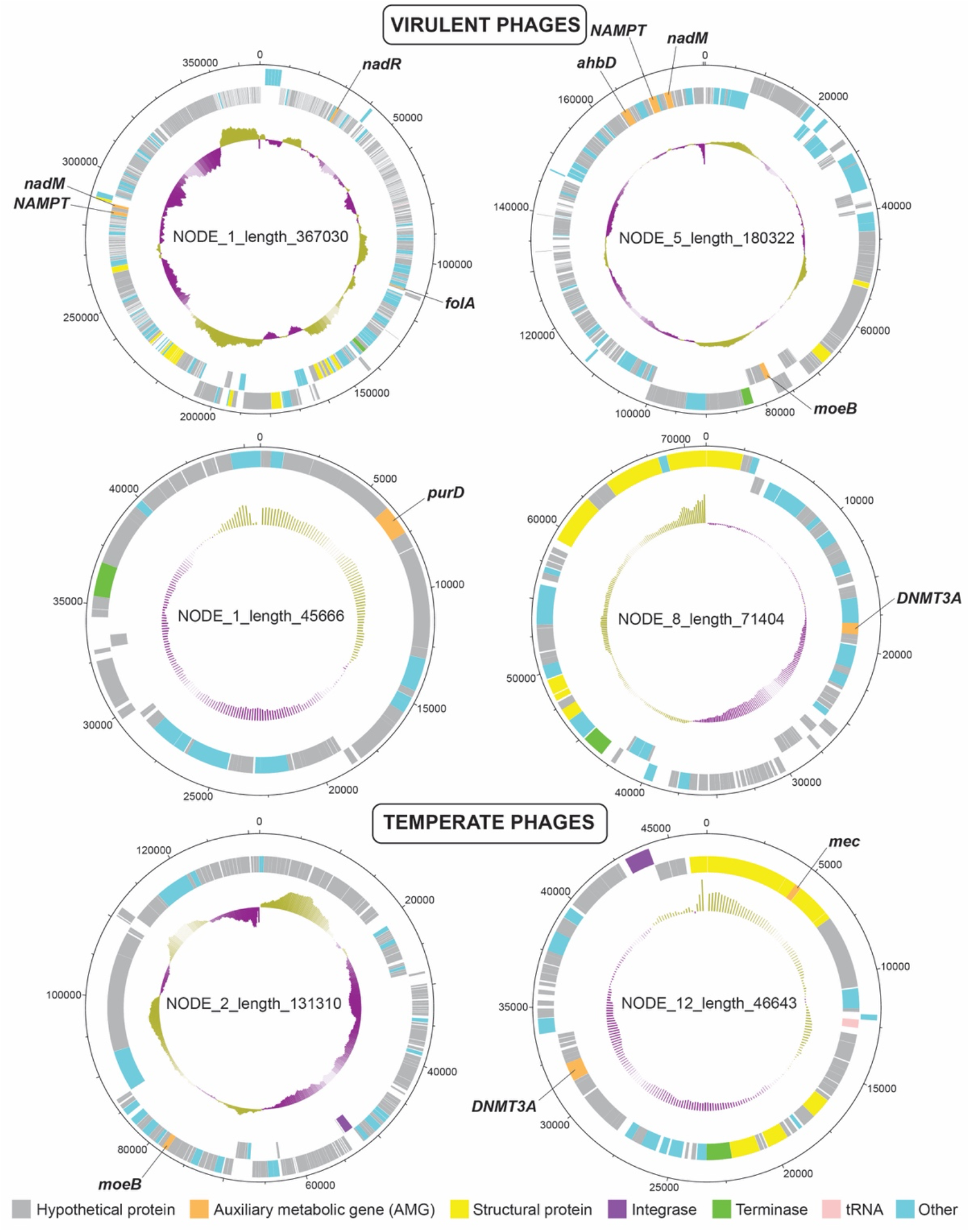
Six complete viral genomes encoding Auxiliary Metabolic Genes (AMGs) in snow. Four of the phages were predicted to be virulent, and two (bottom) were predict
ed to be temperate. The outer ring indicates the nucleotide position in the genome. The two colored middle rings show the predicted genes on the forward strand (outer ring) or reverse strand (inner ring). The innermost ring indicates the GC content of the phage genome. AMGs are annotated. The circular genomes were visualized using DNAplotter (Carver et al., 2009).

## Acknowledgments

The authors would like to acknowledge Heather Maughan for reviewing the manuscript.

## Author Contributions

A.R.B., K.G., F.R., and C.B.S. designed the experiments; A.R.B., K.G., and F.R. performed field sampling and data collection; A.R.B. and C.B.S. generated microscopy, metagenome, and virome data, and performed statistical analyses; A.R.B. wrote the manuscript and all authors made critical revisions; C.B.S. provided guidance, and F.R. provided guidance and funding during the research.

## Declaration of Interests

The authors declare no conflicting interests.

## Data availability

Metagenomic sequences are available in the MG-RAST database under the project name “canada_snow_metagenomes.” Raw virome sequences are available in the NCBI Short Read Archive (SRA) under BioProject ID PRJNA819850.

## Notes

https://www.ncbi.nlm.nih.gov/bioproject/?term=PRJNA819850

https://www.mg-rast.org/mgmain.html?mgpage=project&project=mgp94771

### Competing Interest Statement

The authors have declared no competing interest.

## References

Amato, P., Hennebelle, R., Magand, O., Sancelme, M., Delort, A.-M., Barbante, C., Barbante, C., Boutron, C., & Ferrari, C. (2007). Bacterial characterization of the snow cover at Spitzberg, Svalbard. FEMS Microbiology Ecology, 59(2007), 255–264. doi:10.1111/j.1574-6941.2006.00198.x

Anthenelli, M., Jasien, E., Edwards, R., Bailey, B., Felts, B., Katira, P., Nulton, J., Salamon, P., Rohwer, F., Silveira, C. B., & Luque, A. (2020). Phage and bacteria diversification through a prophage acquisition ratchet. bioRxiv, 2020.04.08.028340. doi:10.1101/2020.04.08.028340

Bondy-Denomy, J. & Davidson, A. R. (2014). When a virus is not a parasite: The beneficial effects of prophages on bacterial fitness. Journal of Microbiology, 52(3), 235–242. doi:10.1007/s12275-014-4083-3

Breitbart, M., Bonnain, C., Malki, K., & Sawaya, N. A. (2018). Phage puppet masters of the marine microbial realm. Nature Microbiology, 3, 754–766. doi:10.1038/s41564-018-0166-y

Breitbart, M., Thompson, L., Suttle, C., & Sullivan, M. (2007). Exploring the Vast Diversity of Marine Viruses. Oceanography, 20(2), 135–139. doi:10.5670/oceanog.2007.58

Bushnell, B. (2014). BBMap: a fast, accurate, splice-aware aligner (No. LBNL-7065E). Lawrence Berkeley National Lab (LBNL), Berkeley, CA (United States).

Canchaya, C., Fournous, G., Chibani-Chennoufi, S., Dillmann, M.-L., & Brüssow, H. (2003). Phage as agents of lateral gene transfer. Current Opinion in Microbiology, 6(4), 417–424. doi:10.1016/S1369-5274(03)00086-9

Carpenter, E. J., Lin, S., & Capone D. G. (2000). Bacterial Activity in South Pole Snow. Applied and Environmental Microbiology, 66(10), 4514–4517. doi:10.1128/AEM.66.10.4514-4517.2000

Carver, T., Thomson, N., Bleasby, A., Berriman, M., & Parkhill, J. (2009). DNAPlotter: circular and linear interactive genome visualization. Bioinformatics, 25(1), 119–120. doi:10.1093/bioinformatics/btn578

De Smet, J., Zimmermann, M., Kogadeeva, M., Ceyssens, P.-J., Vermaelen, W., Blasdel, B., Jang, H. B., Sauer, U., & Lavigne, R. (2016). High coverage metabolomics analysis reveals phage-specific alterations to *Pseudomonas aeruginosa* physiology during infection. ISME Journal, 10(8), 1–13. doi:10.1038/ismej.2016.3

Duval, B., Duval, E., & Hoham, R. W. (1999). Snow algae of the Sierra Nevada, Spain, and High Atlas mountains of Morocco. International Microbiology, 2, 39–42.

Fuhrman, J. A. (1999). Marine viruses and their biogeochemical and ecological effects. Nature, 399(6736), 541–548. doi:10.1038/21119

Fujii, M., Takano, Y., Kojima, H., Hoshino, T., Tanaka, R., & Fukui, M. (2010). Microbial Community Structure, Pigment Composition, and Nitrogen Source of Red Snow in Antarctica. Microbial Ecology, 59, 466–475. doi:10.1007/s00248-009-9594-9

Hampton, H. G., Watson, B. N. J., & Fineran, P. C. (2020). The arms race between bacteria and their phage foes. Nature, 577(7790), 327–336. doi:10.1038/s41586-019-1894-8

Harding, T., Jungblut, A. D., Lovejoy, C., & Vincent, W. F. (2011). Microbes in High Arctic Snow and Implications for the Cold Biosphere. Applied and Environmental Microbiology, 77(10), 3234–3243. doi:10.1128/AEM.02611-10

Hauptmann, A. L., Stibal, M., Bælum, J., Sicheritz-Pontén, T., Brunak, S., Bowman, J. S., Hansen, L. H., Jacobsen, C. S., & Blom, N. (2014). Bacterial diversity in snow on North Pole ice floes. Extremophiles, 18, 945–951. doi:10.1007/s00792-014-0660-y

Hisakawa, N., Quistad, S. D., Hester, E. R., Martynova, D., Maughan, H., Sala, E., Gavrilo, M. V., & Rohwer, F. (2015). Metagenomic and satellite analyses of red snow in the Russian Arctic. PeerJ, 3, 1–13. doi:10.7717/peerj.1491

Hutinet, G., Kot, W., Cui, L., Hillebrand, R., Balamkundu, S., Gnanakalai, S., Neelakandan, R., Carstens, A. B., Lui, C. F., Tremblay, D., Jacobs-Sera, D., Sassanfar, M., Lee, Y., Weigele, P., Moineau, S., Hatfull, G. F., Dedon, P. C., Hansen, L. H., & de Crécy-Lagard, V. (2019). 7-Deazaguanine modifications protect phage DNA from host restriction systems. Nature Communications, 10, 5442. doi:10.1038/s41467-019-13384-y

Kanehisa, M. & Goto, S. (2000). KEGG: Kyoto Encyclopedia of Genes and Genomes. Nucleic Acids Research, 28, 27–30. doi:10.1093/nar/28.1.27

Kieft K., Zhou Z., & Anantharaman K. (2020). VIBRANT: automated recovery, annotation and curation of microbial viruses, and evaluation of virome function from genomic sequences. Microbiome, 8, 90. doi:10.1186/s40168-020-00867-0

Knowles, B., Silveira, C. B., Bailey, B. A., Barott, K., Cantu, V. A.,;Cobián-Güemes, A. G., Coutinho, F. H., Dinsdale, E. A., Felts, B., Furby, K. A., George, E. E., Green, K. T., Gregoracci, G. B., Haas, A. F., Haggerty. J. M., Hester, E. R., Hisakawa, N., Kelly, L. W., Lim, Y. W.,… Rohwer, F. (2016). Lytic to temperate switching of viral communities. Nature, 531(7595), 466–470. doi:10.1038/nature17193

Kozloff, L. M., Lute, M., & Crosby, L. K. (1977). Bacteriophage T4 Virion Baseplate Thymidylate Synthetase and Dihydrofolate Reductase. Journal of Virology, 23(3), 637–644. doi:10.1128/jvi.23.3.637-644.1977

Krug, L., Erlacher, A., Markut, K., Berg, G., & Cernava, T. (2020). The microbiome of alpine snow algae shows a specific inter-kingdom connectivity and algae-bacteria interactions with supportive capacities. ISME Journal, 14(9), 2197–2210. doi:10.1038/s41396-020-0677-4

Krüger, D. H. & Bickle, T. A. (1983). Bacteriophage Survival: Multiple Mechanisms for Avoiding the Deoxyribonucleic Acid Restriction Systems of Their Hosts. Microbiological Reviews, 47 (3), 345–360. doi:10.1128/mmbr.47.3.345-360.1983

Lau, C., Niere, M., & Ziegler, M. (2009). The NMN/NaMN adenylyltransferase (NMNAT) protein family. Frontiers in Bioscience, 14(2), 410–431. doi:10.2741/3252

Lee, J. Y., Li, Z., & Miller, E. S. (2017). Vibrio phage KVP40 Encodes a Functional NAD+ Salvage Pathway. Journal of Bacteriology, 199(9), e00855–16. doi:10.1128/JB.00855-16

Leonardi, R., Zhang, Y. M., Rock, C. O., & Jackowski, S. (2005). Coenzyme A: Back in action. Progress in Lipid Research, 44, 125–153. doi:10.1016/j.plipres.2005.04.001

Li, W. & Godzik, A. (2006). Cd-hit: A fast program for clustering and comparing large sets of protein or nucleotide sequences. Bioinformatics, 22(13), 1658–1659. doi:10.1093/bioinformatics/btl158

Lim, Y. W., Haynes, M., Furlan, M., Robertson, C. E., Harris, J. K., & Rohwer, F. (2014). Purifying the Impure: Sequencing Metagenomes and Metatranscriptomes from Complex Animal-associated Samples. Journal of Visualized Experiments, (94), e52117. doi:10.3791/52117

Liu, Y., Yao, T., Jiao, N., Kang, S., Xu, B., Zeng, Y., Huang, S., & Liu, X. (2009). Bacterial diversity in the snow over Tibetan Plateau Glaciers. Extremophiles, 13, 411–423. doi:10.1007/s00792-009-0227-5

Lopatina, A., Krylenkov, V., & Severinov, K. (2013). Activity and bacterial diversity of snow around Russian Antarctic stations. Research in Microbiology, 164(9), 949–958. doi:10.1016/j.resmic.2013.08.005

Luo, E., Eppley, J. M., Romano, A. E., Mende, D. R., & DeLong, E. F. (2020). Double-stranded DNA virioplankton dynamics and reproductive strategies in the oligotrophic open ocean water column. ISME Journal, 14, 1304–1312. doi:10.1038/s41396-020-0604-8

Luque, A. & Silveira, C. B. (2020). Quantification of Lysogeny Caused by Phage Coinfections in Microbial Communities from Biophysical Principles. mSystems, 5(5), e00353–20. doi:10.1128/mSystems.00353-20

Lutz, S., Anesio, A. M., Raiswell, R., Edwards, A., Newton, R. J., Gill, F., & Benning, L. G. (2016). The biogeography of red snow microbiomes and their role in melting arctic glaciers. Nature Communications, 7, 11968. doi:10.1038/ncomms11968

Maccario, L., Vogel, T. M., & Larose, C. (2014). Potential drivers of microbial community structure and function in Arctic spring snow. Frontiers in Microbiology, 5, 413. doi:10.3389/fmicb.2014.00413

Mačková, M., Boháčová, S., Perlíková, P., PoštováSlavětínská, L., & Hocek, M. (2015). Polymerase Synthesis and Restriction Enzyme Cleavage of DNA Containing 7-Substituted 7-Deazaguanine Nucleobases. ChemBioChem, 16 (15), 2225–2236. doi:10.1002/cbic.201500315

Mathews, C. K. (1967). Growth of a Dihydrofolate Reductaseless Mutant of Bacteriophage T4. Journal of Virology, 1(5), 963–967. doi:10.1128/jvi.1.5.963-967.1967

McNair, K., Bailey, B. A., & Edwards, R. A. (2012). PHACTS, a computational approach to classifying the lifestyle of phages. Bioinformatics, 28, 614–618. doi:10.1093/bioinformatics/bts014

Meyer, F., Paarmann, D., D’Souza, M., Olson, R., Glass, E. M., Kubal, M., Paczian, T., Rodriguez, A., Stevens, R., Wilke, A., Wilkening, J., & Edwards, R. A. (2008). The metagenomics RAST server – a public resource for the automatic phylogenetic and functional analysis of metagenomes. BMC Bioinformatics, 9, 386. doi:10.1186/1471-2105-9-386

Michaud, L., Lo Giudice, A., Mysara, M., Monsieurs, P., Raffa, C., Leys, N., Amalfitano, S., & Van Houdt, R. (2014). Snow Surface Microbiome on the High Antarctic Plateau (DOME C). PLoS ONE, 9(8), e104505. doi:10.1371/journal.pone.0104505

Murat Eren, A., Esen, O. C., Quince, C., Vineis, J. H., Morrison, H. G., Sogin, M. L., & Delmont, T. O. (2015). Anvi’o: An advanced analysis and visualization platform for’omics data. PeerJ, 2015(10), 1–29. doi:10.7717/peerj.1319

Murphy, J., Mahony, J., Ainsworth, S., Nauta, A., & Sinderen, V. (2013). Bacteriophage Orphan DNA Methyltransferases: Insights from Their Bacterial Origin, Function, and Occurrence. Applied and Environmental Microbiology, 79(24), 7547–7555. doi:10.1128/AEM.02229-13

Nishimura, Y., Yoshida, T., Kuronishi, M., Uehara, H., Ogata, H., & Goto, S. (2017). ViPTree: The viral proteomic tree server. Bioinformatics, 33(15), 2379–2380. doi:10.1093/bioinformatics/btx157

Nurk, S., Meleshko, D., Korobeynikov, A., & Pevzner, P. A. (2017). MetaSPAdes: A new versatile metagenomic assembler. Genome Research, 27(5), 824–834. doi:10.1101/gr.213959.116

Phillips, G., El Yacoubi, B., Lyons, B., Alvarez, S., Iwata-Reuyl, D., & de Crécy-Lagard, V. (2008). Biosynthesis of 7-Deazaguanosine-Modified tRNA Nucleosides: A New Role for GTP Cyclohydrolase I. Journal of Bacteriology, 190(24), 7876–7884. doi:10.1128/JB.00874-08

Ponstingl, H. & Ning, Z. (2010). SMALT - A New Mapper for DNA Sequencing Reads. F1000Posters, 1(313). Retrieved from https://f1000research.com/posters/327 %0A http://cdn.f1000.com/posters/docs/327

Puxty, R. J., Evans, D. J., Millard, A. D., & Scanlan, D. J. (2018). Energy limitation of cyanophage development: Implications for marine carbon cycling. ISME Journal, 12(5), 1273–1286. doi:10.1038/s41396-017-0043-3

Reader, J. S., Metzgar, D., Schimmel, P., & de Crécy-Lagard, V. (2004). Identification of Four Genes Necessary for Biosynthesis of the Modified Nucleoside Queuosine. Journal of Biological Chemistry, 279(8), 6280–6285. doi:10.1074/jbc.M310858200

Revelle, W. (2021). psych: Procedures for Psychological, Psychometric, and Personality Research. Northwestern University, Evanston, Illinois. R package version 2.1.6, https://CRAN.R-project.org/package=psych.

Revollo, J. R., Grimm, A. A., & Imai, S. I. (2007). The regulation of nicotinamide adenine dinucleotide biosynthesis by Nampt/PBEF/visfatin in mammals. Current Opinion in Gastroenterology, 23(2), 164–170. doi:10.1097/MOG.0b013e32801b3c8f

Rohwer F., Segall, A., Steward, G., Seguritan, V., Breitbart, M., Wolven, F., & Azam, F. (2000). The complete genomic sequence of the marine phage Roseophage SIO1 shares homology with nonmarine phages. Limnology and Oceanography, 45(2), 408–418. doi: 10.4319/lo.2000.45.2.0408

Samson, J. E., Magadán, A. H., Sabri, M., & Moineau, S. (2013). Revenge of the phages: Defeating bacterial defences. Nature Reviews Microbiology, 11(10), 675–687. doi:10.1038/nrmicro3096

Sánchez-Romero, M. A., Cota, I., & Casadesús, J. (2015). DNA methylation in bacteria: From the methyl group to the methylome. Current Opinion in Microbiology, 25, 9–16. doi:10.1016/j.mib.2015.03.004

Segawa, T., Miyamoto, K., Ushida, K., Agata, K., Okada, N., & Kohshima, S. (2005). Seasonal change in bacterial flora and biomass in mountain snow from the Tateyama Mountains, Japan, analyzed by 16S rRNA gene sequencing and real-time PCR. Applied and Environmental Microbiology, 71(1), 123–130. doi:10.1128/AEM.71.1.123-130.2005

Silveira, C. B., Luque, A., Roach, T. N. F., Villela, H., Barno, A., Green, K., Reyes, B., Rubio-Portillo, E., Le, T., Mead, S., Hatay, M., Vermeij, M. J. A., Takeshita, Y., Haas, A., Bailey, B., & Rohwer, R. (2019). Biophysical and physiological processes causing oxygen loss from coral reefs. eLife, 8, e49114. doi:10.7554/eLife.49114

Silveira, C. B., Luque, A., & Rohwer, F. (2021). The landscape of lysogeny across microbial community density, diversity and energetics. Environmental Microbiology, 00, 1–14. doi:10.1111/1462-2920.15640

Simon, C., Wiezer, A., Strittmatter, A. W., & Daniel, R. (2009). Phylogenetic Diversity and Metabolic Potential Revealed in a Glacier Ice Metagenome. Applied and Environmental Microbiology, 75(23), 7519–7526. doi:10.1128/AEM.00946-09

Sorci, L., Martynowski, D., Rodionov, D. A., Eyobo, Y., Zogaj, X., Klose, K. E., Nikolaev, E. V., Magni, G., Zhang H., & Osterman, A. L. (2009). Nicotinamide mononucleotide synthetase is the key enzyme for an alternative route of NAD biosynthesis in Francisella tularensis. PNAS, 106(9), 3083–3088. doi:10.1073/pnas.0811718106

Suttle, C. A. (2005). Viruses in the sea. Nature, 437(7057), 356–361. doi:10.1038/nature04160

Terashima, M., Umezawa, K., Mori, S., Kojima, H., & Fukui, M. (2017). Microbial community analysis of colored snow from an alpine snowfield in Northern Japan reveals the prevalence of Betaproteobacteria with snow algae. Frontiers in Microbiology, 8, 1481. doi:10.3389/fmicb.2017.01481

Thingstad, T. F. (2000). Elements of a theory for the mechanisms controlling abundance, diversity, and biogeochemical role of lytic bacterial viruses in aquatic systems. Limnology and Oceanography, 45(6), 1320–1328. doi:10.4319/lo.2000.45.6.1320

Thomas, W. H. & Duval, B. (1995). Sierra Nevada, California, U.S.A., Snow Algae: Snow Albedo Changes, Algal-Bacterial Interrelationships, and Ultraviolet Radiation Effects. Arctic and Alpine Research, 27(4), 389–399. doi:10.1080/00040851.1995.12003136

Thompson, L. R., Zeng, Q., Kelly, L., Huang, K. H., Singer, A. U., Stubbe, J., & Chisholm, S. W. (2011). Phage auxiliary metabolic genes and the redirection of cyanobacterial host carbon metabolism. PNAS, 108(39), E757-E764. doi:10.1073/pnas.1102164108

Thurber, R. V., Haynes, M., Breitbart, M., Wegley, L., & Rohwer, F. (2009). Laboratory procedures to generate viral metagenomes. Nature Protocols, 4(4), 470–483. doi: 10.1038/nprot.2009.10

Tock, M. R. & Dryden, D. T. F. (2005). The biology of restriction and anti-restriction. Current Opinion in Microbiology, 8(4), 466–472. doi:10.1016/j.mib.2005.06.003

Vinayak, M. & Pathak, C. (2010). Queuosine modification of tRNA: its divergent role in cellular machinery. Bioscience Reports, 30(2), 135–148. doi:10.1042/bsr20090057

Walker, J. E. (1992). The NADH: ubiquinone oxidoreductase (complex I) of respiratory chains. Quarterly Reviews of Biophysics, 25(3), 253–324. doi:10.1017/S003358350000425X

Weiss, R. L. (1983). Fine Structure of the Snow Alga *(Chlamydomonas nivalis)* and Associated Bacteria. Journal of Phycology, 19, 200–204. doi:10.1111/j.0022-3646.1983.00200.x

Yakimovich, K. M., Engstrom, C. B., & Quarmby, L. M. (2020). Alpine Snow Algae Microbiome Diversity in the Coast Range of British Columbia. Frontiers in Microbiology, 11, 1721. doi:10.3389/fmicb.2020.01721

Zeng, Q. & Chisholm, S. W. (2012). Marine viruses exploit their host’s two-component regulatory system in response to resource limitation. Current Biology, 22(2), 124–128. doi:10.1016/j.cub.2011.11.055

