## Supplementary Figure for "Viral and bacterial ecogenomics in globally expanding red snow blooms"

**Supplementary Figures:**

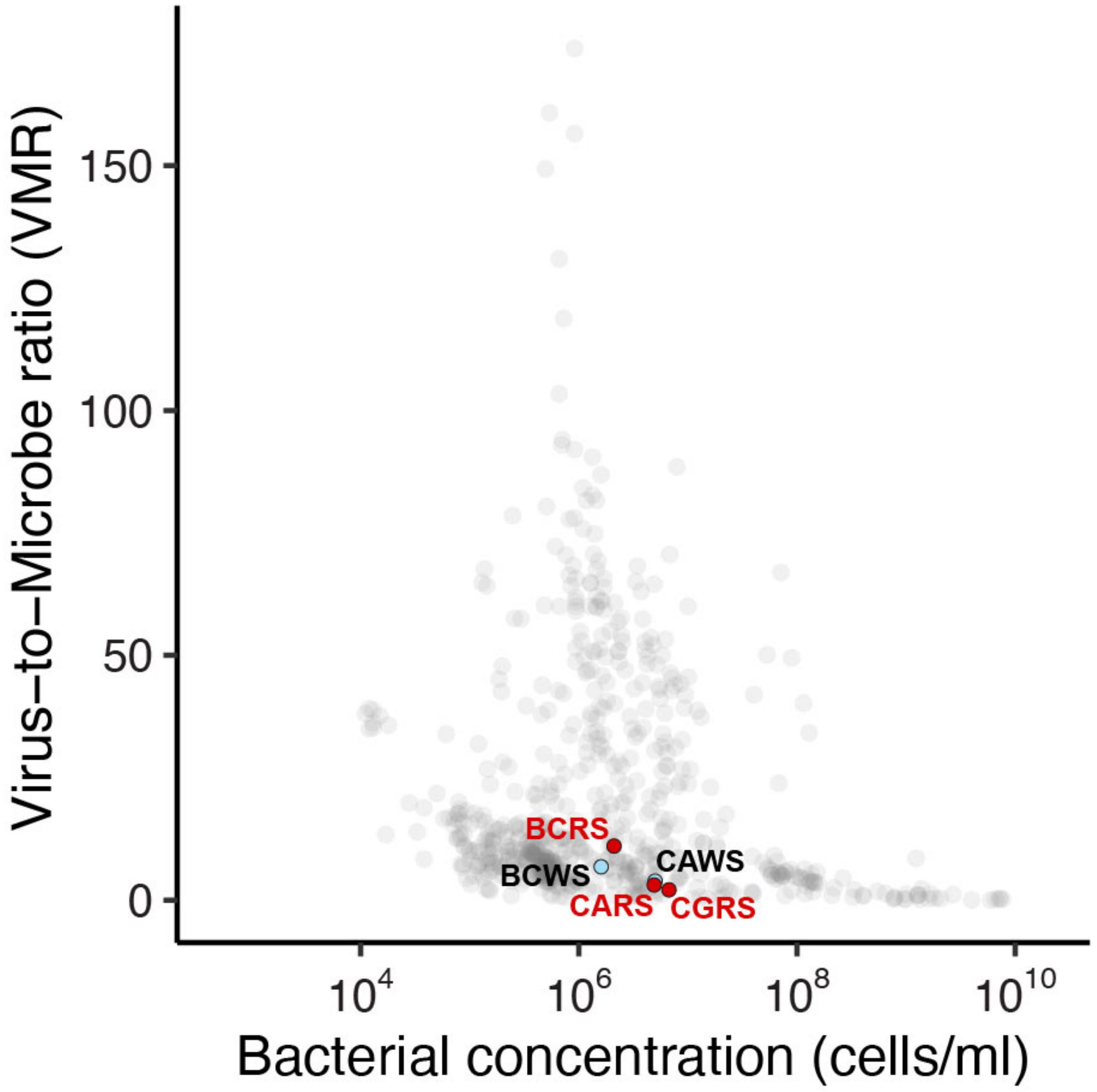


Supplementary Figure 1. Virus-to-microbe ratios (VMRs) of the five snow samples plotted with VMRs from marine, freshwater, soil, and animal-associated ecosystems (from Knowles et al., 2016). Cell abundances, virus counts, and VMR data from the snow samples are in Table 2. BCWS = Blackcomb Mountain White Snow, BCRS = Blackcomb Mountain Red Snow, CAWS = Callaghan Pass White Snow, CARS = Callaghan Pass Red Snow, CGRS = Cougar Mountain Red Snow.

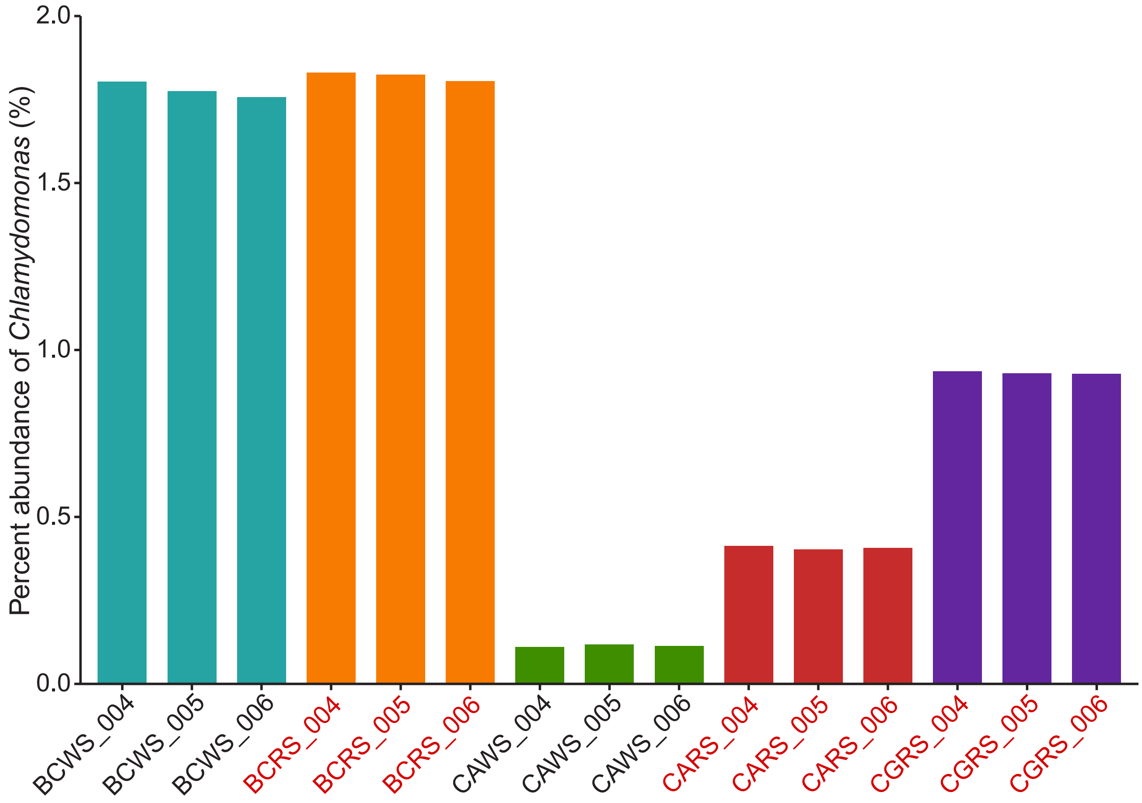


Supplementary Figure 2. Percent abundance of the algae genus *Chlamydomonas* in each metagenome, as classified by MG-RAST searchers against the M5NR database. BCWS = Blackcomb Mountain White Snow, BCRS = Blackcomb Mountain Red Snow, CAWS = Callaghan Pass White Snow, CARS = Callaghan Pass Red Snow, CGRS = Cougar Mountain Red Snow.

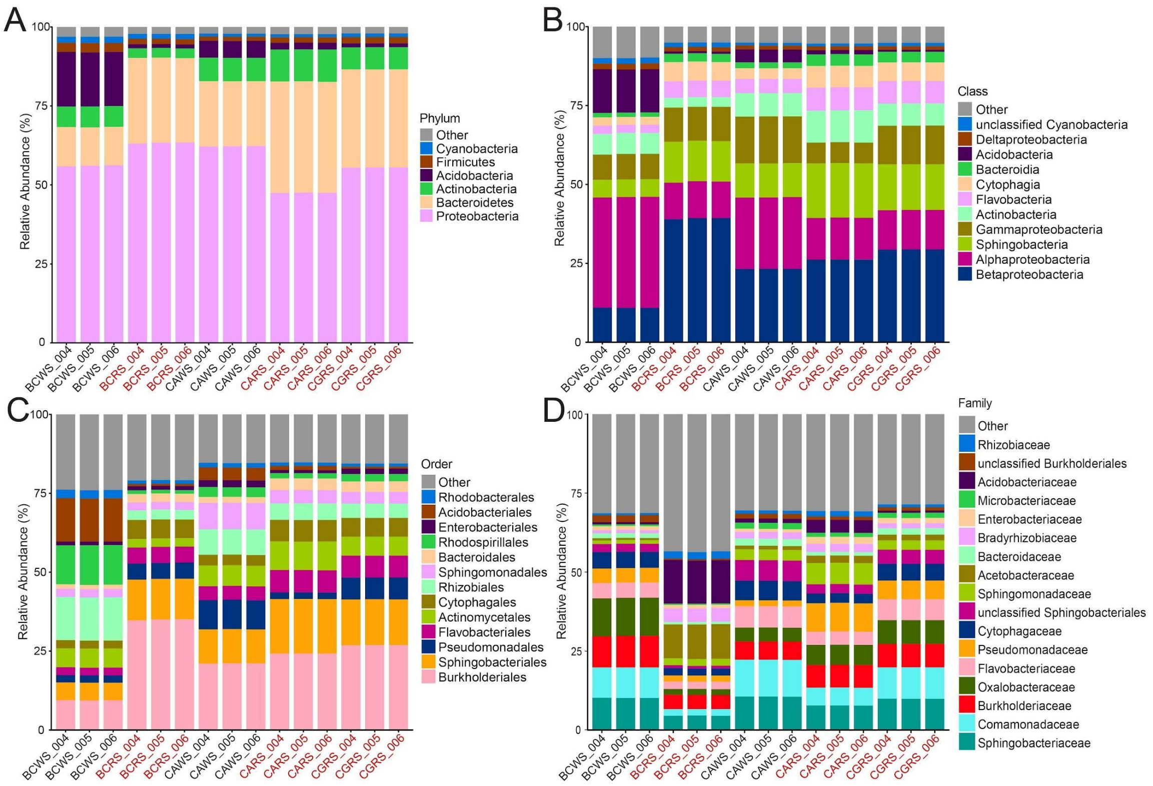


Supplementary Figure 3. Bacterial taxonomic profiles. The plots display taxa comprising over 1% of the total bacterial reads. (A) phylum, (B) class, (C) order, and (D) family. BCWS = Blackcomb Mountain White Snow, BCRS = Blackcomb Mountain Red Snow, CAWS = Callaghan Pass White Snow, CARS = Callaghan Pass Red Snow, CGRS = Cougar Mountain Red Snow.

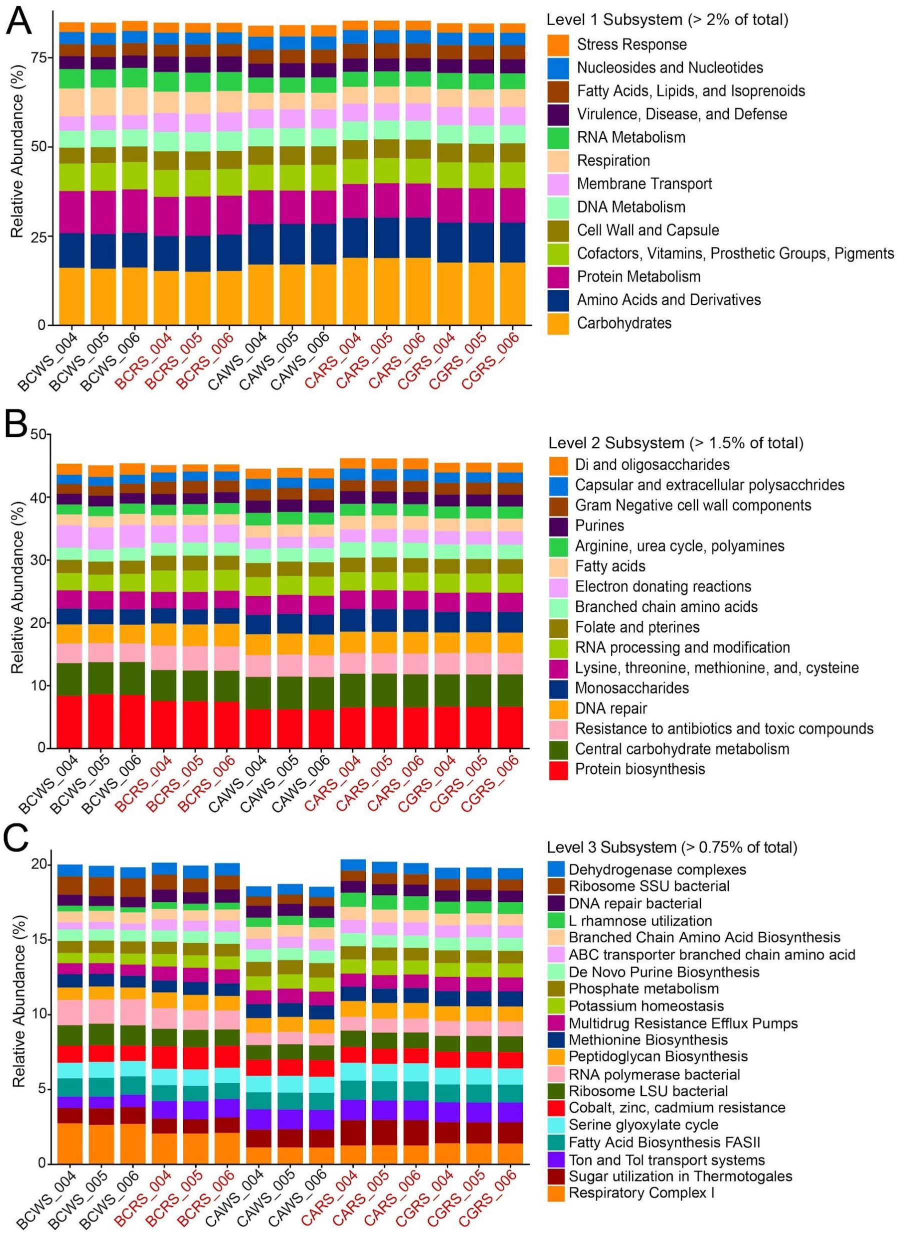


Supplementary Figure 4. The SEED level subsystems for the Canada snow metagenomes. (A) SEED level 1, (B) SEED level 2, and (C) SEED level 3. BCWS = Blackcomb Mountain White Snow, BCRS = Blackcomb Mountain Red Snow, CAWS = Callaghan Pass White Snow, CARS = Callaghan Pass Red Snow, CGRS = Cougar Mountain Red Snow.
